# Assessment of Repurposed Compounds for Antiviral Activity Against Measles Virus

**DOI:** 10.64898/2026.03.31.715719

**Authors:** Annika Rössler, Jose Ayala-Bernot, Shadi Mohammadabadi, Ninaad Lasrado, Siddhesh Warke, Robert Flaumenhaft, Dan H. Barouch

## Abstract

**Background:** There is currently no approved antiviral therapy against measles virus (MeV). Repurposing available compounds with broad antiviral activity may rapidly identify candidate drugs for clinical evaluation. Here we evaluated the antiviral activity of the clinically approved drugs azelastine hydrochloride and zafirlukast as well as the flavonoids quercetin and isoquercetin against MeV in preventative and therapeutic *in vitro* studies.

**Methods:** Compounds were tested for antiviral activity against MeV in preventative (prophylactic and virucidal) and therapeutic (steady-state and persistent) assays in Vero/hSLAM cells. Viral loads and cell viability were measured 48h post-infection, and dose-response curves were used to calculate EC50 values. Flavonoids were also tested in the presence of 1 mM ascorbic acid.

**Results:** Azelastine hydrochloride did not show evidence of antiviral activity against MeV under these conditions, whereas zafirlukast, quercetin, and isoquercetin showed therapeutic activity against MeV. The addition of ascorbic acid enhanced the therapeutic potency of quercetin to 4.2-4.8 µM and of isoquercetin to 10.7-10.9 µM. Antiviral activity was dose-dependent when administered post-infection.

**Conclusion:** Among the four compounds tested, quercetin showed the most potent therapeutic antiviral activity against MeV *in vitro*. Isoquercetin and zafirkulast also showed therapeutic activity. These findings support further evaluation of quercetin, isoquercetin, and zafirlukast as candidate antiviral drugs for MeV and highlight the utility of *in vitro* platforms for rapid antiviral drug screening.

## Introduction

Measles virus (MeV) is one of the most contagious human pathogens with a high reproductive ratio (R_0_) of 12 to 18 and can cause severe complications such as pneumonia, encephalitis, and death, particularly in vulnerable populations (1). MMR vaccine coverage of >95% has been shown to maintain population immunity and limit outbreaks (1-3). Although MeV was declared eliminated in the United States in 2000, defined as the absence of continuous MeV transmission for one year in the presence of surveillance programs (4), the epidemiologic landscape is now characterized by recurrent outbreaks (2). In 2024, 285 confirmed MeV cases were reported in the US, which increased to 2,283 in 2025, and more than 1,500 cases have been reported this year by March 2026, with 92% of cases in individuals who were unvaccinated or had an unknown vaccine status (5).

The two dose MMR vaccine provides up to 97% protection (6), but effective countermeasures are still needed for breakthrough cases and for outbreak settings. Moreover, immunocompromised individuals may respond poorly to the vaccine. At present, no approved therapy for MeV infection is available, and disease management is largely limited to supportive care (7). This represents an unmet medical need given the increasing MeV outbreaks (5) and the decreasing vaccine rates (8).

Repurposing compounds that have broad antiviral activity and that are either clinically available or in clinical trials may identify potential candidate drugs against MeV that could be advanced rapidly into clinical trials. Azelastine hydrochloride, zafirlukast, and the flavonoids quercetin and its derivative isoquercetin have shown broad antiviral activity, but their activity against MeV has not yet been evaluated. Here we evaluated the antiviral activity of these four compounds against MeV using *in vitro* preventive and therapeutic screening approaches.

## Methods

### Cell line

Vero/hSLAM cells (Cat. 04091501; MilliporeSigma) stably expressing human signaling lymphocytic activation molecule (SLAM, CD150) were cultured in DMEM supplemented with 5% FBS and maintained under selection with 0.4 mg/mL Geneticin to preserve the expression of the CD150 receptor. Cells were maintained in humidified incubators at 37 °C containing 5% CO_2_ atmosphere.

### Measles virus

The MeV isolate (MVs/Ohio.USA/17.14/3 [D9]) was obtained through BEI Resources (NR-52252) and propagated on Vero/hSLAM cells.

### Virus stock production

For all MeV infection experiments, DMEM supplemented with 2% FBS was used. Vero/hSLAM cells were seeded the day before infection and infected with MeV at an MOI of 0.01 at approximately 90% confluency. One day after infection, the supernatant (SN) was replaced by fresh medium. Cells were inspected daily for syncytia formation and once cytopathic effect appeared (usually 3 days post infection) tissue culture flasks were frozen at -80 °C. Tissue culture flasks were thawed at 37 °C to release cell-associated MeV particles and SN was clarified by centrifugation (2,400 rpm, 5 min, RT). Subsequently, clarified SN was underlaid with 30% sucrose and virus was purified via centrifugation (29,000 x g, 2h, 4 °C). The SN was carefully aspirated and the virus containing pellet was recovered in pre-cooled PBS and kept for 15 min on ice, before virus aliquots were prepared and cryopreserved at -80 °C.

### Virus titration

The titers of MeV stocks were determined by a TCID_50_ assay. Vero/hSLAM cells were seeded in 96-well plates with 1×10^5^ cells/well the day before infection. Half-log dilution series of MeV stocks were prepared in DMEM containing 2% FBS and added to the cells. CPE was evaluated 5 days post infection and the TCID_50_ titer was determined according to the Spearman-Kärber method.

### Antiviral compounds

The antiviral compounds tested included azelastine hydrochloride (Cat. A7611; Millipore-Sigma, USA), zafirlukast (Cat. Z4152; Millipore-Sigma, USA), quercetin (Cat. Q4951; Millipore-Sigma, USA), and isoquercetin (quercetin-3-β-D-glucoside; Cat. 17793; Millipore-Sigma, USA). All compounds were reconstituted in sterile DMSO, and single-use aliquots were stored at -80C.

### Antiviral screening against MeV

Vero/hSLAM cells were seeded in 96-well plates with 1×10^5^ cells/well the day before infection. The antiviral compounds were screened for antiviral capacity against MeV in duplicates in two preventive (prophylactic and virucidal) and two therapeutic (steady-state and persistent) settings. Across all experimental settings, compounds were prepared as 2-fold serial dilutions in media (DMEM supplemented with 2% FBS or DMEM supplemented with 2% FBS and 1 mM ascorbic acid) and tested at final concentrations ranging from 50 to 3.13 µM. The purified MeV stock was diluted to infect cells at an MOI of 0.01 across all experiments. For all experiments, two studies were run in parallel in 96-well plates to allow assessment of cell viability and viral loads.

#### Prophylactic setting

The SN of cells was aspirated and diluted compounds were added and incubated with the cells for 1h at 37 °C. Afterwards, the antiviral compounds were aspirated, and MeV was added to the cells. Cells and virus were incubated for 1h at 37 °C before SN was aspirated, cells were washed and fresh media was added. Cells were incubated for 48h at 37 °C prior readout.

#### Virucidal setting

Diluted compounds and MeV virus were mixed at a 1:1 ratio and preincubated for 1h at 37 °C. The SN of cells was aspirated and the antiviral/virus mixture was transferred to the cells and incubated for 1h at 37 °C. Subsequently, the SN was aspirated, cells were washed and fresh media was added. Cells were incubated for 48h at 37 °C prior readout.

#### Steady-state and persistent settings

The SN of cells was aspirated and cells were infected with MeV for 1h at 37 °C. The virus was aspirated, cells were washed and diluted compounds were added to the cells. For both therapeutic settings, cells were incubated for a total of 48h after infection at 37 °C prior readout. To maintain the drug concentration over time in the steady-state setting, the SN was aspirated and replaced by freshly diluted compounds 24h post infection. No re-dosing was performed in the persistent setting.

### Cell Viability

Cell viability was assessed using a commercially available luminescence-based cell viability assay (CellTiter-Glo® 2.0 Cell Viability Assay, Cat. G9241, Promega, USA), following a protocol adapted from the manufacturer’s instructions. Briefly, SN of cells was carefully aspirated, PBS and CellTiter-Glo® substrate were added at a 1:1 ratio. Tissue plates were incubated on an orbital shaker for 10 mins at RT prior luminescence measurement. 100% viability was assumed in medium control wells (without MeV infection), while cells incubated with 1% Triton-X/PBS for 1h prior viability assessment served as negative control (0% viability).

### Viral Load Quantification

#### Sample Preparation

Two days after infection, tissue culture plates were frozen at -80 °C and thawed at RT. The culture SN was then mixed and lysed in a 1:1 ratio with modified DLR buffer for isolation of viral RNA(9), containing 0.5% IGEPAL, 25mM NaCl in 10 mM Tris-HCl buffer, RNA carrier (Qiagen, Germany) and Ribonuclease inhibitor (Cat. 9134, Aldevron, USA).

#### Quantitative RT-PCR

Viral RNA was quantified by qRT-PCR targeting the MeV hemagglutinin gene (MeV-H). A standard was generated by subcloning the target gene into a pcDNA3.1(+) expression plasmid, using restriction site cloning (Integrated DNA Technologies). The insert was subsequently *in-vitro* transcribed into RNA using the HiScribe T7 High Yield RNA Synthesis Kit (New England Biolabs, E2040S). Log dilutions of the standard were prepared for RT-PCR assays ranging from 1E10 copies to 1E−1 copies.

Standard dilutions and extracted RNA of samples were reverse transcribed into cDNA using SuperScript VILO III Master Mix (Invitrogen, 11755050) according to the manufactuers instructions. Quantitative PCR was performed using TaqMan Fast Advanced Master Mix (Life Technologies, 4444965) on QuantStudio 6 and 7 Flex Real-Time PCR Systems (Applied Biosystems). The assay sequences are as follows:

MeV-H.Fwd: CAATCGAGCATCAGGTCAAGG;

MeV-H.Rev: GTCCTCAGGCCCACTTCATC;

MeV-H.Probe:/56-FAM/CTCGCAAAG/ZEN/GCGGTTACGGCC/3IABkFQ/.

Thermal cycling conditions consisted of an initial denaturation step at 95 °C for 20s, followed by 45 cycles of 95 °C for 1s and 60° C for 20s, with annealing, extension, and fluorescence detection performed in a single step. Standard curves were generated for each run to calculate MeV-H RNA copies per milliliter. The limit of detection was 250 copies/mL.

## Data Analysis

Viral loads were normalized to untreated virus-only control wells which were set to 100%, with the limit of detection (LOD; 250 copies/mL) set to 0%. Cell viability was normalized to medium-only control wells which were set to 100%, with 0% viability defined by the relative luminescence units (RLU) obtained from maximal cell death control wells treated with 1% Triton X-100 in PBS. For each compound, concentration, and experimental setting, duplicate measurements were first averaged within each independent experiment, and summary data of two or more independent experiments are shown as mean ± standard error of the mean (SEM) across biological replicates. Dose-response curves were fitted using a four-parameter log-logistic (4PL) model implemented in the drc package (v3.0-1) in R (v4.5.0)(10-12). EC50 values were estimated from the fitted curves.

## Results

All compounds were screened in four experimental settings, referred to as prophylactic, virucidal, steady-state, and persistent settings, to evaluate preventative and therapeutic antiviral activity against MeV at different stages of infection. In the virucidal setting, MeV was preincubated with the compounds before infection of susceptible Vero/hSLAM cells to examine direct effects on viral particles. In the prophylactic setting, cells were pre-treated with antivirals prior infection to evaluate interference with early stages of infection. Therapeutic activity was assessed in MeV-infected cells with daily compound replenishment (steady-state) or with single dose treatment (persistent). For all experiments, cell viability and viral loads were measured 48h post infection. The results with azelastine hydrochloride and zafirlukast are shown in **Figure 1**, and the results with the flavonoids quercetin and isoquercetin are shown in **Figure 2** and **Figure 3**.

**Figure 1.**
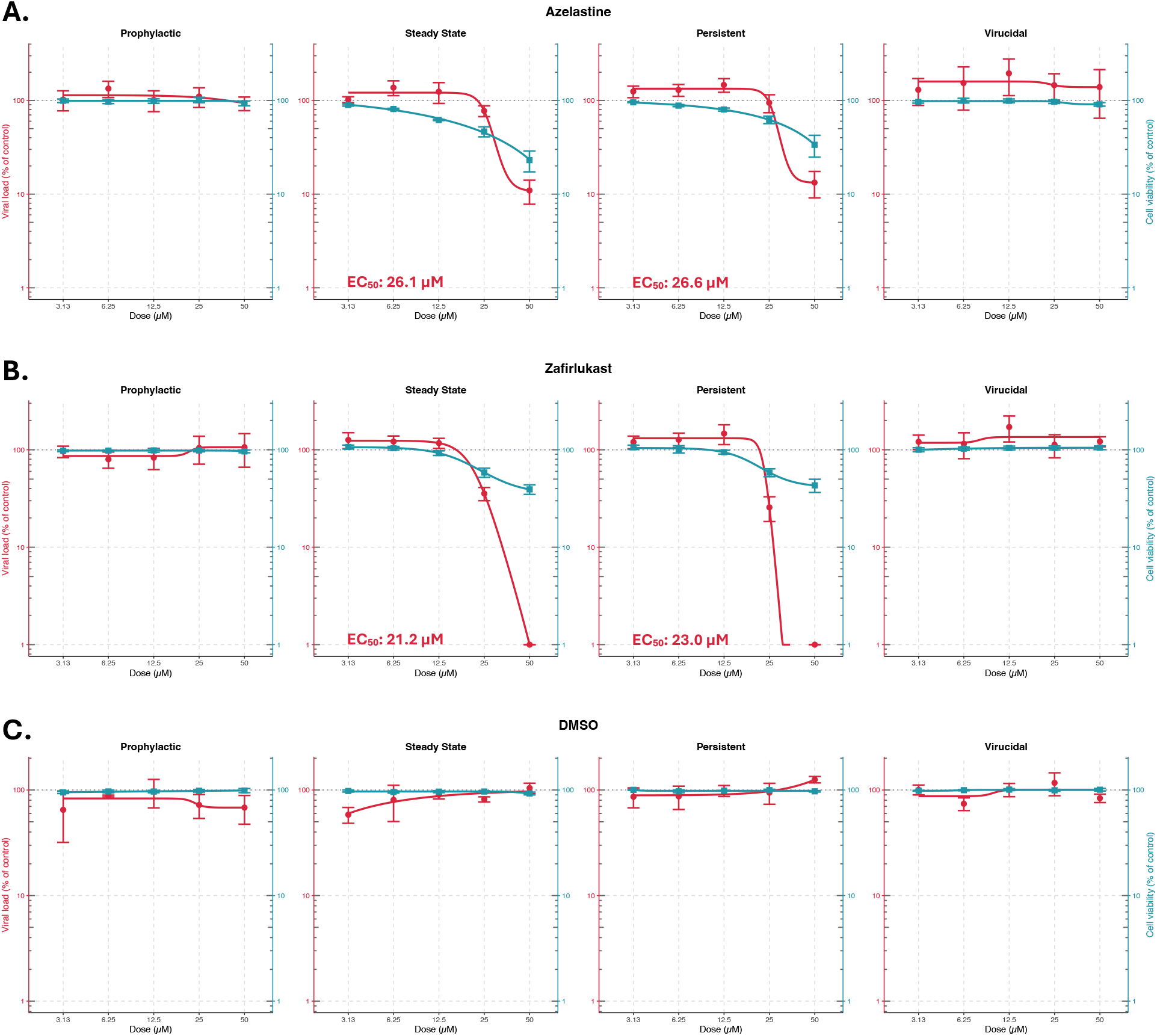
Antiviral efficacy of azelastine hydrochloride and zafirlukast against MeV across four experimental settings. Azelastine hydrochloride **(A)**, zafirlukast **(B)**, and the DMSO control **(C)** were titrated and tested for antiviral activity against MeV in prophylactic, steady-state, persistent, and virucidal settings, and viral loads and cell viability were analyzed 48h post infection. Viral loads were normalized to virus-only controls (100%) with the limit of detection (LOD; 250 copies/mL) set to 0%. Cell viability was assessed using CellTiter-Glo® assay (Promega) and normalized to the medium-only control (100%), with 0% viability defined by a maximal cell death control generated by incubating cells with 1% Triton X-100/PBS for 1h prior readout. Data points represent mean ± SEM of independent replicates and solid lines indicate interpolated 4-parameter logistic (4PL) curve, with viral loads shown in red and viability represented in blue. EC50 values for viral loads are shown in red when an accurate and biologically relevant curve fit for viral load data was obtained.

**Figure 2.**
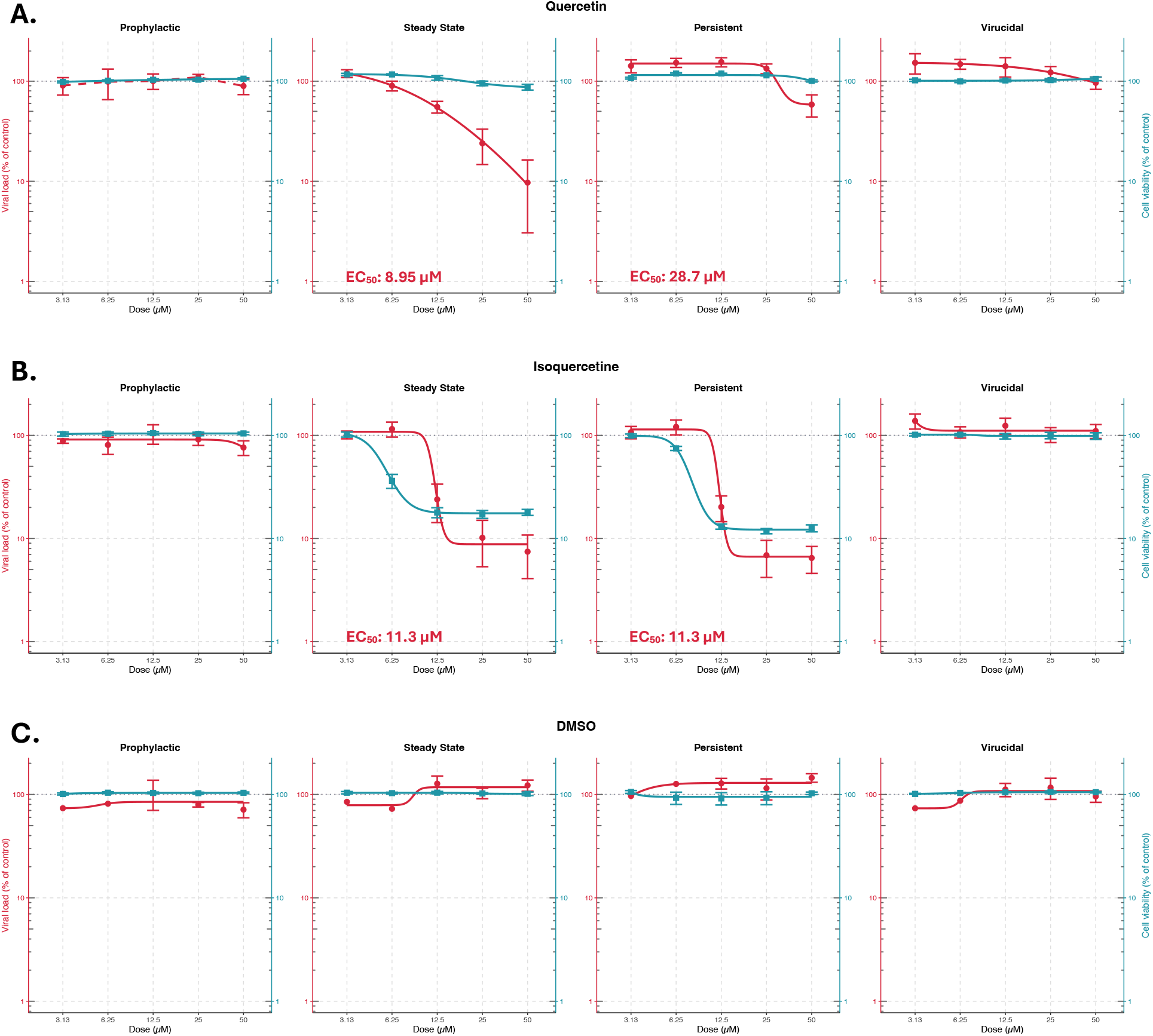
Antiviral efficacy of flavonoids against MeV in four experimental settings. Quercetin **(A)** and isoquercetin **(B)** were tested against MeV in prophylactic, steady-state, persistent, and virucidal settings, with DMSO as the control **(C)**. Viral loads and cell viability were evaluated 48h post infection. Viral loads were normalized to virus-only controls (100%) with the limit of detection (LOD; 250 copies/mL) set to 0%. Cell viability was assessed using CellTiter-Glo® assay (Promega) and normalized to the medium-only control (100%), with 0% viability defined by a maximal cell death control generated by incubating cells with 1% Triton X-100/PBS for 1h prior readout. Data points represent mean ± SEM of independent replicates and solid lines indicate interpolated 4-parameter logistic (4PL) curve, with viral loads shown in red and viability represented in blue. EC50 values for viral loads are shown in red when an accurate and biologically relevant curve fit for viral load data was obtained.

**Figure 3.**
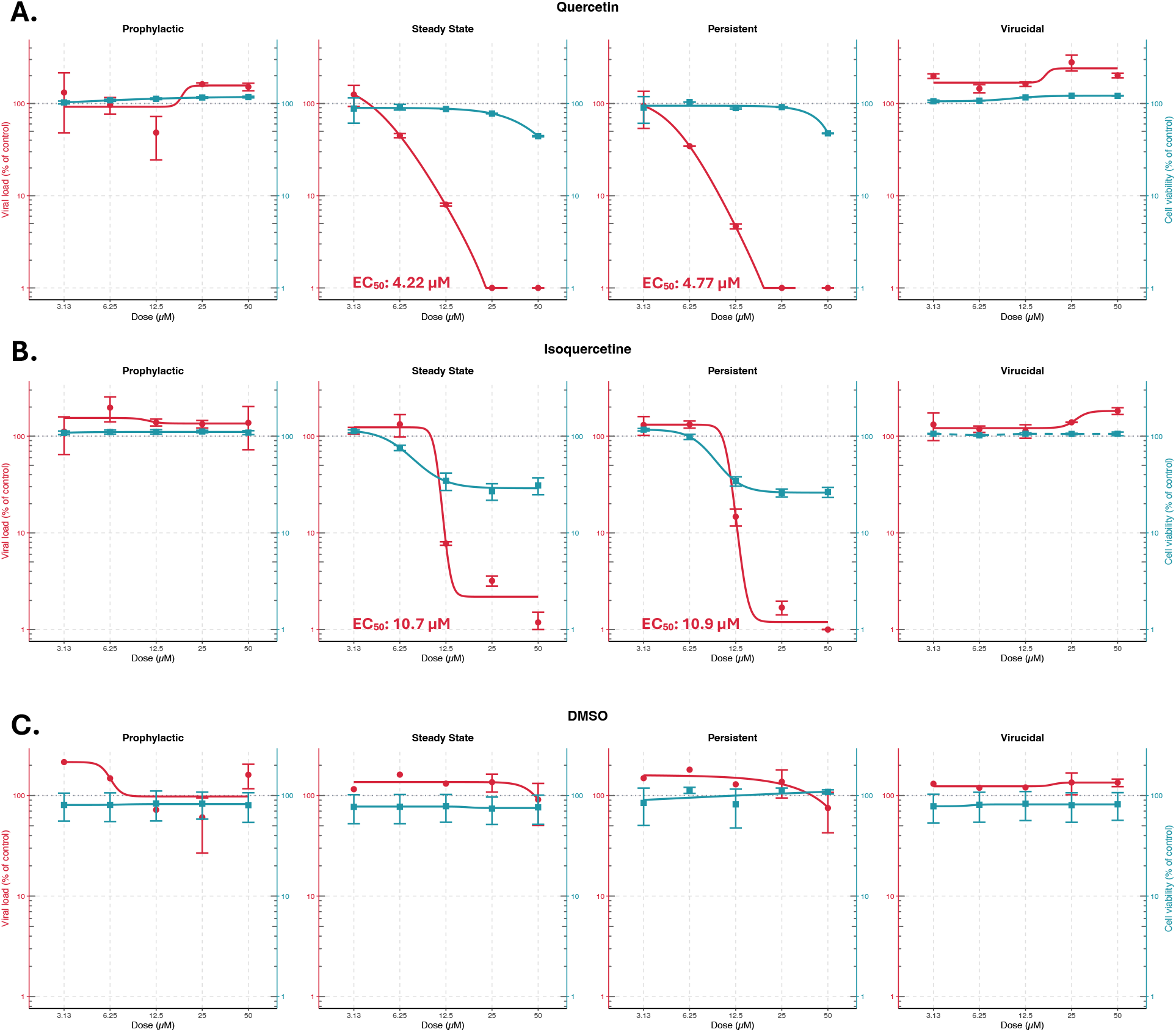
Antiviral efficacy of flavonoids against MeV in presence of 1 mM ascorbic acid across four experimental settings. Quercetin **(A)** and isoquercetin **(B)** in medium supplemented with 1 mM ascorbic acid were tested against MeV in prophylactic, steady-state, persistent, and virucidal settings, with DMSO as the control **(C)**. Viral loads and cell viability were evaluated 48h post infection. Viral loads were normalized to virus-only controls (100%) with the limit of detection (LOD; 250 copies/mL) set to 0%. Cell viability was assessed using CellTiter-Glo® assay (Promega) and normalized to the medium-only control (100%), with 0% viability defined by a maximal cell death control generated by incubating cells with 1% Triton X-100/PBS for 1h prior readout. Data points represent mean ± SEM of independent replicates and solid lines indicate interpolated 4-parameter logistic (4PL) curve, with viral loads shown in red and viability represented in blue. EC50 values for viral loads are shown in red when an accurate and biologically relevant curve fit for viral load data was obtained.

Azelastine hydrochloride did not show clear antiviral activity against MeV in the tested conditions (**Fig. 1A**). Although viral loads decreased at higher concentrations of azelastine, cell viability declined to a comparable extent, which reduced the sensitivity of the assay to detect antiviral activity. In contrast, zafirlukast showed clear dose-dependent antiviral activity in the therapeutic experiments (**Fig. 1B**), with a half-maximal effective concentration (EC_50_) of 21.2 µM in the steady-state (multiple-dose) setting and 23.0 µM in the persistent (single-dose) setting. No substantial antiviral activity was observed in the prophylactic or virucidal settings for these compounds.

Quercetin (**Fig. 2A**) showed clear dose-dependent antiviral activity against MeV in the steady-state setting with an EC_50_ value of 8.95 µM but less activity in the persistent setting. We observed similar trends for isoquercetin, but cell viability decreased proportionately (**Fig. 2B**). Since flavonoids undergo rapid oxidation in cell culture media that can lead to the generation of hydrogen peroxide that may affect cell viability (13-15), we repeated these experiments with the flavonoids in the presence of 1 mM ascorbic acid as an antioxidant. Under these conditions, we observed enhanced antiviral potency for both quercetin (**Fig. 3A**) and isoquercetin (**Fig. 3B**), although the activity of quercetin remained higher and isoquercetin led to some reduced cell viability. The EC_50_ values for quercetin were 4.22 µM and 4.77 µM and the EC_50_ values for isoquercetin were 10.7 µM and 10.9 µM for the steady-state and persistent therapeutic settings, respectively. No antiviral activity was observed with DMSO controls (**Figs. 1C, 2C, 3C**).

## Discussion

Despite vaccination and public health surveillance efforts, MeV outbreaks continue to occur, and current treatment is limited to supportive care. In this study, we evaluated four compounds with broad antiviral activity for potential antiviral activity against MeV in preventative and therapeutic *in vitro* settings. Azelastine hydrochloride is an approved antihistamine, zafirkulast is an approved leukotriene antagonist, and quercetin and isoquercetin are flavonoids in clinical development for other conditions that are designed by the FDA as Generally Recognized As Safe (GRAS). We observed dose-dependent antiviral therapeutic effects for zafirlukast, quercetin, and isoquercetin, particularly in the presence of ascorbic acid.

*In vitro* studies have shown broad antiviral activity of azelastine against multiple respiratory viruses, including SARS-CoV-2, HCoV-229E and RSV (16). In contrast, we did not observe antiviral activity of azelastine against MeV, although the sensitivity of these assays to detect MeV activity was limited by reduced cell viability. For SARS-CoV-2, the antiviral mechanism has been linked to the inhibition of ACE-2 dependent cell entry (16). Azelastine has recently shown protective activity against SARS-CoV-2 and other respiratory viruses in humans in a phase 2 randomized clinical trial (17, 18). In contrast, MeV enters cells by receptor binding via H protein and membrane fusion is mediated by the F protein at the cell surface (19).

Zafirlukast has been reported to inhibit ZIKV and DENV by disruption of viral particles (20). Although we did not observe direct virucidal activity against MeV, zafirlukast showed post-entry therapeutic activity against MeV with EC_50_ values of 21–23 µM, which is comparable with its antiviral potency against SARS-CoV-2 in Vero cells of 25 µM (21) and its potency against West Nile Virus of 32 µM (22).

Quercetin and its derivatives have shown broad antiviral activity by interfering with viral attachment and entry, genome replication, protein syntheses, particle assembly, and host immune responses (23, 24). Quercetin has been reported to have antiviral activity against multiple respiratory viruses, including influenza virus, SARS-CoV-2, and rhinovirus (RV), as well as against other viruses, such as HSV, ZIKV, and DENV (23-29). Its efficacy against influenza virus was mainly linked to inhibition of infection, but its efficacy against SARS-CoV-2 and RV was associated with altered virus replication dynamics (25, 27). Quercetin exhibited antiviral effects against ZIKV in virucidal and treatment settings with EC_50_ values of 11.9 µM and 28.8 µM, respectively (28). In our studies, quercetin demonstrated antiviral activity against MeV in therapeutic settings with improved potency in presence of ascorbic acid with EC_50_ values of 4.2 and 4.8 µM in the steady-state and persistent settings, respectively. We did not observe antiviral activity of quercetin against MeV in the prophylactic and virucidal settings, potentially due to short pre-incubation times of the experiment (28).

The enhanced antiviral activity of quercetin and isoquercetin observed with ascorbic acid likely reflects its antioxidant effects on stabilization of the flavonoids in tissue culture media (13-15). Isoquercetin, which is a glycosylated derivate of quercetin, showed antiviral activity against MeV in the presence of ascorbic acid. Its activity appeared lower than that of quercetin, potentially reflecting its reduced lipophilic properties *in vitro*. Nevertheless, isoquercetin may offer significant advantages *in vivo* due to its improved gastrointestinal absorption and bioavailability following oral administration (24, 30, 31). Recent data also suggest that isoquercetin and zafirlukast may exhibit enhanced biological activity in combination (32).

This study has several important limitations. First, antiviral activity was evaluated *in vitro* in experiments in Vero/hSLAM cells, which provide a practical and high throughput screening platform but do not capture the complexity of MeV infection *in vivo*. Second, antiviral effects were assessed by monitoring viral loads and cell viability rather than clinical disease endpoints. Third, certain compounds such as azelastine and isoquercetin reduced cell viability *in vitro*, which limited the sensitivity of these assays to detect antiviral activity. Fourth, the mechanism of the antiviral activity of these compounds against MeV remains to be determined.

Our data show antiviral therapeutic activity against MeV *in vitro* for several repurposed compounds with broad antiviral activity. Of the four compounds tested, quercetin showed the most robust antiviral activity, followed by zafirkulast and isoquercetin. These findings suggest that further evaluation of these compounds is warranted both *in vitro* and *in vivo*. Our studies also highlight the utility of the *in vitro* platforms for rapid initial screening of candidate antiviral therapeutic drugs.

## Acknowledgements

We thank Daniel Drexler, Thomas Lines, Daniel Kennedy, and William Archer for generous advice, assistance, and reagents. We acknowledge BARDA/VITAL/Start2 75A50124C00012.2025-DEV-1.01 and philanthropic grants (D.H.B.) for supporting this work.

